# Endogenous activity modulates stimulus and circuit-specific neural tuning and perception

**DOI:** 10.1101/687152

**Authors:** Yuanning Li, Michael J. Ward, R. Mark Richardson, Max G’Sell, Avniel Singh Ghuman

**Affiliations:** Center for the Neural Basis of Cognition, Carnegie Mellon University and University of Pittsburgh, Pittsburgh, PA; Program in Neural Computation and Machine Learning, Carnegie Mellon University and University of Pittsburgh, Pittsburgh, PA; Department of Neurological Surgery, University of Pittsburgh, Pittsburgh, PA; Department of Statistics, Carnegie Mellon University, Pittsburgh, PA

**Author notes:** Correspondence to: Yuanning Li. Department of Neurological Surgery, University of California, San Francisco, CA.

## Abstract

Perception reflects not only input from the sensory periphery, but also the endogenous neural state when sensory inputs enter the brain. Whether endogenous neural states influence perception only through global mechanisms, such as arousal, or can also perception in a neural circuit and stimulus specific manner remains largely unknown. Intracranial recordings from 30 pre-surgical epilepsy patients showed that endogenous activity independently modulated the strength of trial-by-trial neural tuning of different visual category-selective neural circuits. Furthermore, the same aspect of the endogenous activity that influenced tuning in a particular neural circuit also correlated with reaction time only for trials with the category of image that circuit was selective for. These results suggest that endogenous activity may influence neural tuning and perception through circuit-specific predictive coding processes.

## Main Text

Perception depends on not only sensory input, but also the neural and cognitive state when a stimulus is presented. Traditionally, this endogenous activity has been treated as random biological noise(*1*). However, studies in both humans and animals demonstrate that rather than being a noise process, endogenous activity reflects fluctuations of neural activity that influence neural processing in a behaviorally relevant manner. Specifically, endogenous fluctuations in neural activity influence both coarse aspects of the neural response to sensory input(*2-5*) and the behavioral response to that input(*5-9*). Endogenous activity has rich structure, reflecting the stimulus processing properties of the local neural circuitry(*10*), broad scale brain network architecture(*11*), and may reflect statistically optimal representations of the environment(*12*). Fuctuations in endogenous processes such as arousal(*13-15*) and alertness(*16, 17*) can influence stimulus processing and behavior. Some theoretical accounts posit that fluctuations of endogenous activity can reflect predictive processes(*18*) that facilitate stimulus processing in a stimulus-specific manner. However, most studies have only examined nonspecific mechanisms, such as arousal and alertness(*13-17*). Thus, there is a dearth of empirical evidence testing whether endogenous processes can influence neural tuning and influence behavior in a circuit and stimulus-specific manner as required by models of predictive processing.

Data were acquired from 30 human neurosurgical patients with implanted intracranial electroencephalography (iEEG) while they viewed images of faces, bodies, words, hammers, houses, and scrambled non-objects and performed a 1-back, repeat detection task. Stimuli were balanced across categories and presented in a random order to reduce any potential cognitive or strategic processes that might favor one stimulus over another. This allowed us to probe the relationship between endogenous activity, visual category tuning, and behavior separately for different categories of stimuli and separately for the cortical circuits selective to these categories without bias. Analyses identified 246 iEEG electrodes selective for one of these visual categories that were then used for the primary analyses examining the effects of endogenous activity on category selectivity. These iEEG electrodes were distributed across the cortex, though concentrated in the bilateral ventral temporal cortex (VTC) (Fig. 1, Table 1). Three main hypotheses were tested sequentially: 1. endogenous activity modulates the strength of category tuning in response to visual stimuli; 2. the same aspect of the endogenous activity that modulates tuning also correlates with behavioral perception in a region-by-stimulus specific manner, where endogenous activity in regions selective for a particular stimulus will only correlate with behavior for that stimulus (e.g. endogenous activity in regions selective for faces will correlate with behavioral performance for face stimuli); 3. the aspect of endogenous activity that modulates tuning and behavior is uncorrelated across regions selective for different visual categories. Support for these three hypotheses would suggest that endogenous fluctuations can modulate stimulus-specific visual category tuning, the same aspect of the activity that modulates tuning also influences behavior, and that this modulation does not reflect an unspecific phenomenon, such as arousal, but rather differentially and independently influences circuits selective for different categories of visual stimuli. Additional analyses elucidated further details about what aspects of the endogenous activity modulate stimulus-specific category tuning and behavior.

**Table 1.**
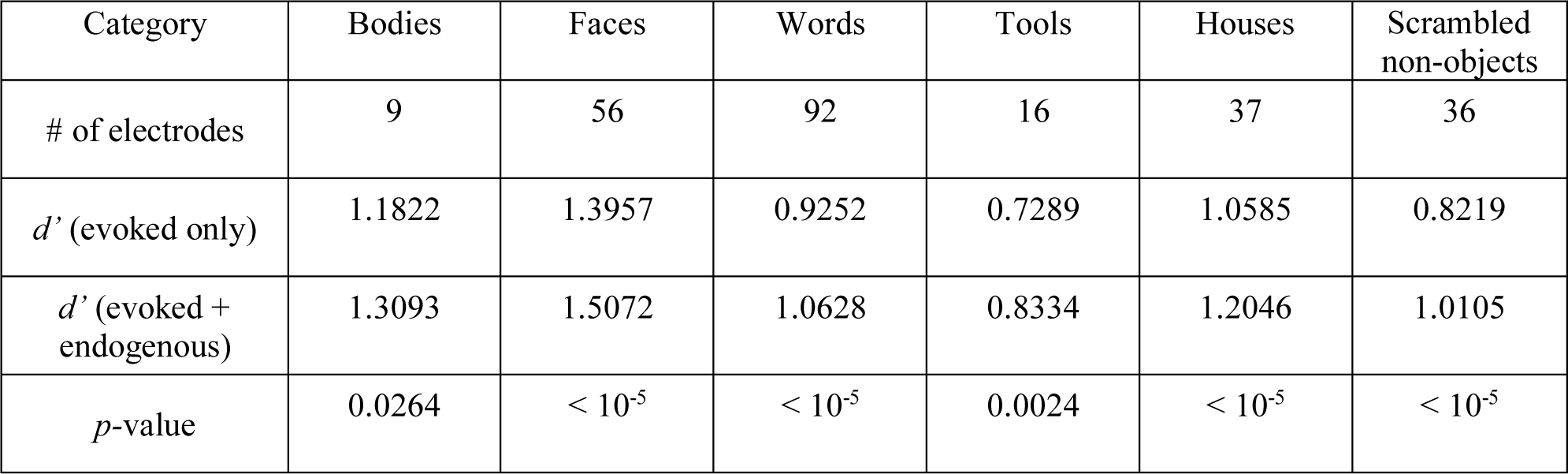
Number of electrodes showing significant category sensitivity for each of the stimulus categories, and the comparisons of classification results from the two-stage GLM

**Fig. 1.**
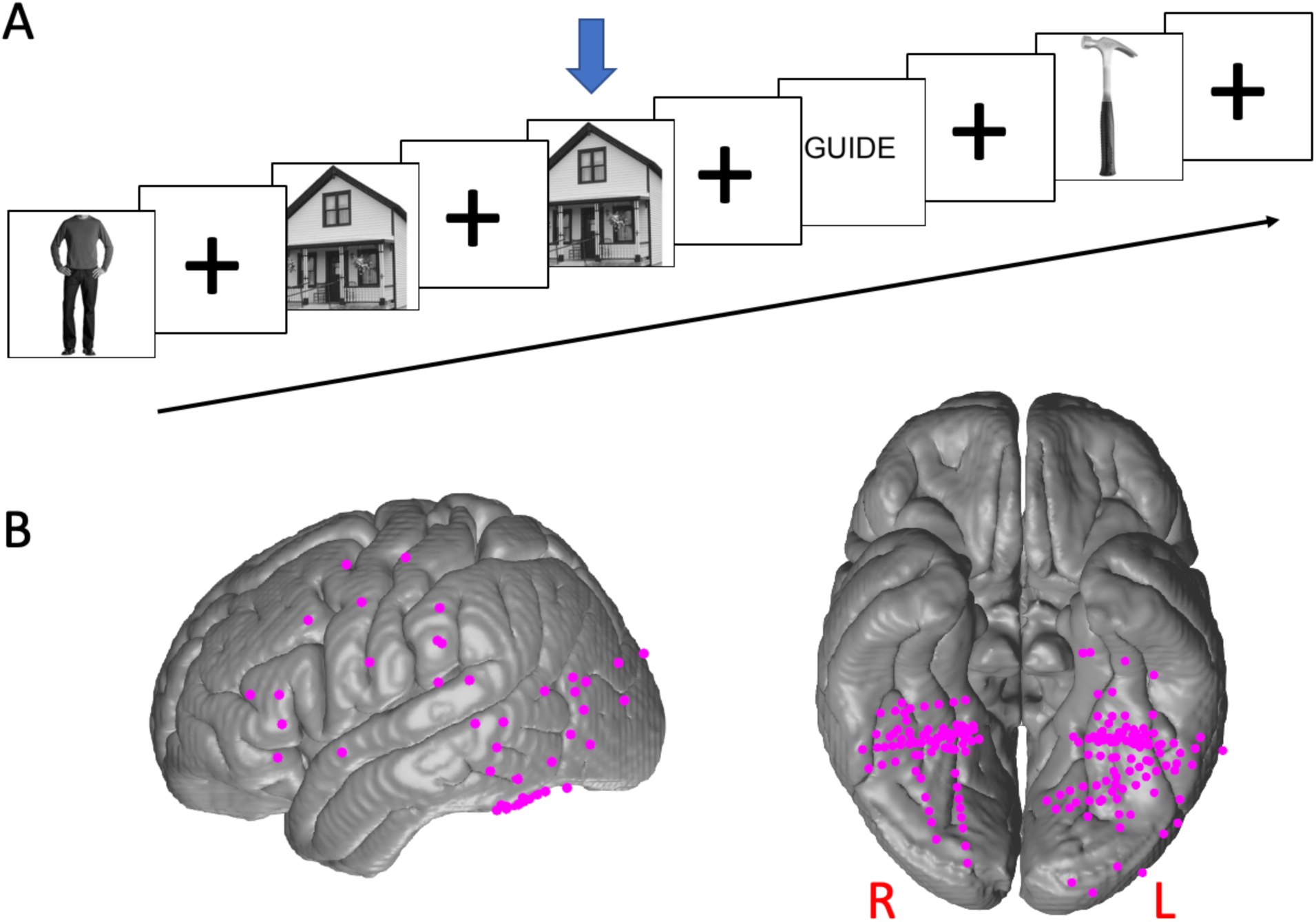
Behavioral task and the localization of the category-selective electrodes. **A)** Experimental paradigm in which the subject is shown a series of images and performs a 1-back repeat detection task. 180 images from 6 categories (faces, bodies, words, tools, houses, scrambled non-objects) were used. Each image was presented for 900 ms, with 900 ms interstimulus interval. **B)** The left lateral and bilateral ventral views of the locations of the 246 category-selective electrodes mapped onto a common brain surface. The category-selectivity was determined based on 1) significant sensitivity index (*d’*) for certain category using a 6-way classifier; 2) larger event-related potential (mean stFP) or mean stHFA over other categories.

Previous studies in humans have established the relationship between features of endogenous activity and the evoked response, including oscillatory phase and power/amplitude of the event-related response (*19-21*) or blood oxygen-level dependent (BOLD) signal (*22*). While these studies show that endogenous activity may affect the stimulus evoked response, they do not establish whether it can modulate the quality of the neural representation for stimuli in ways that are related to perception, such as the strength of tuning for particular stimuli. A common way to study population-level neural tuning is to use a multivariate discriminant model to assess the separability of the population neural activity with regard to different categories(*23, 24*). Specifically, discriminant models extract critical dimensions in the space of evoked response that discriminate the preferred category from the others (*24*).

To evaluate the modulation effect along the critical discriminant dimensions, pre-stimulus activity, including single-trial field potential (stFP), single-trial broadband high-frequency activity (stBHA), and phases at different frequencies, was used as a proxy for the endogenous neural state of the brain at the time of stimulus presentation. Specifically, a model was used to modulate classification boundary along the critical discriminating directions, based on the pre-stimulus activity, and the resulting improvement in accuracy was examined. Because the pre-stimulus activity contains no information about the conditions (see supplemental results), the only way classification accuracy can be improved using this model is if the pre-stimulus activity contains information about how strongly tuned the stimulus response along the critical dimensions will be on a particular trial.

The algorithm is designed to use this pre-stimulus information, if it is present, to adjust the classification boundary, i.e. trial-by-trial tuning, in the discriminant dimension on each trial to optimize classification. Comparing classification accuracy with and without this adjustment tests the first hypothesis that endogenous activity modulates neural tuning. In addition, this adjustment provides a trial-by-trial measure of how much influence endogenous activity has on neural tuning, which we term the “modulation index” (MI). The correlation between MI and behavioral reaction time on a simple perceptual task tests the second hypothesis that the same aspect of the endogenous activity that modulates tuning also correlates with behavioral perception. Comparing the correlation of the MI between pairs of electrodes that record from areas selective for the same versus different categories of visual stimuli test the third hypothesis that the aspect of the endogenous activity that modulates tuning is stimulus-specific and thus uncorrelated across circuits selective for different stimuli.

The results indicated that conditioning the model on pre-stimulus activity improved the classification accuracy for all visual categories, compared to the classification accuracy using only post-stimulus activity (Fig. 2A, Table 1). One potential confound is the pre-stimulus activity could reflect cognition related to the previous trial and particularly repetitions of the same condition. For example, if subjects were presented two face trials in a row, the pre-stimulus activity for the second trial could reflect lingering activity from the first trial. This potential confound was addressed by demonstrating that classification accuracy improves with inclusion of the pre-stimulus activity, even after accounting for trial order effects, particularly repetitions of the same condition (Table S1). These results show that critical features of pre-stimulus activity relate to the strength of neural tuning and that modifying the discriminant model based on this relationship improves classification accuracy. Therefore, these results support the first hypothesis that endogenous activity modulates the degree of category tuning in response to visual stimuli.

**Fig. 2.**
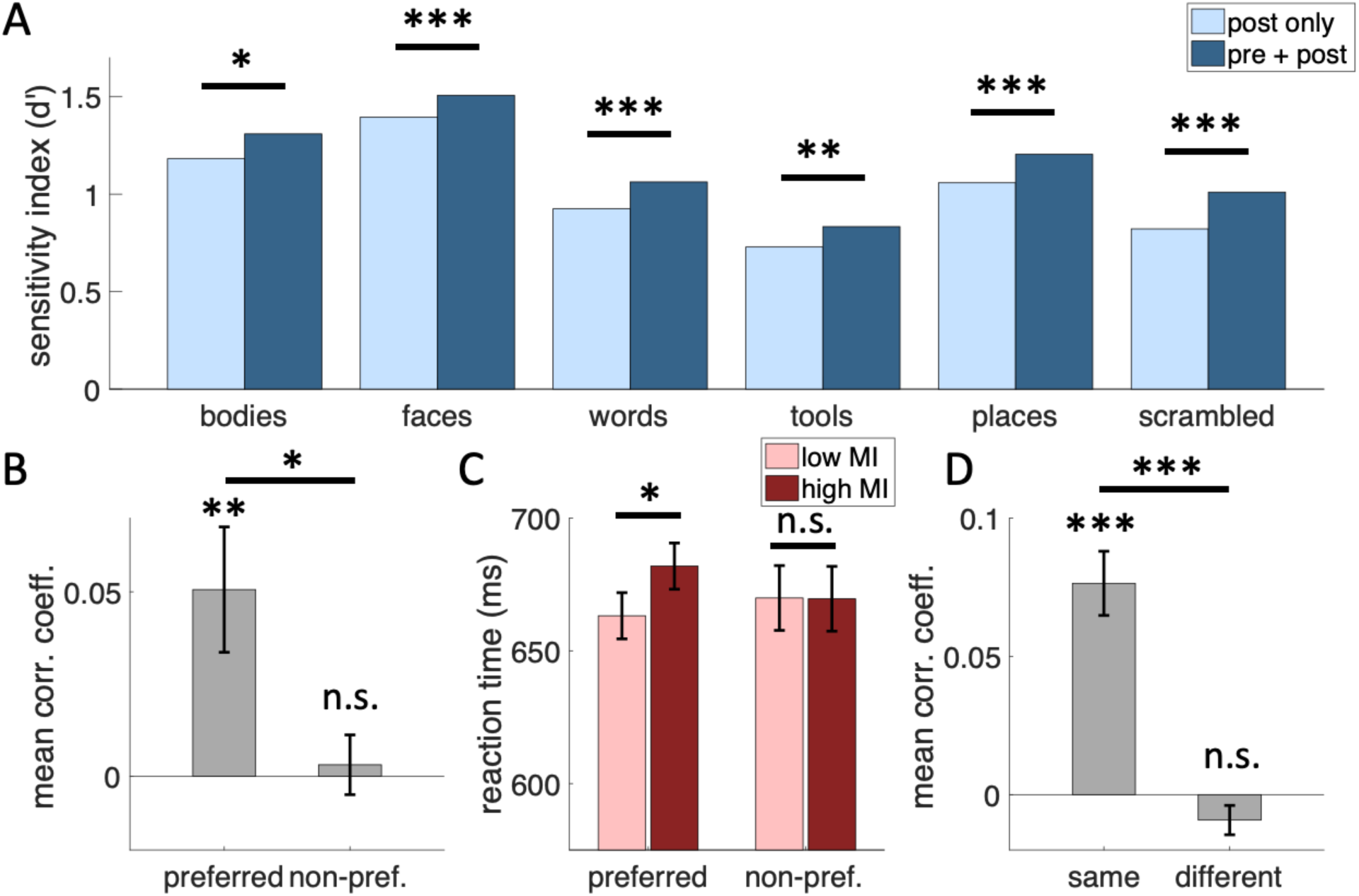
Pre-stimulus endogenous activity influenced post-stimulus category tuning and perceptual behavior in a neural circuit and stimulus specific manner. **A)** Category classification accuracy (mean sensitivity index *d*’) before (post only) and after (pre + post) conditioning on endogenous activity. Mean *d*’ = 1.06 without conditioning on pre-stimulus activity versus mean *d*’ = 1.19 after conditioning on pre-stimulus activity (t(245)= 12.39; *p* < 1×10^−5^, paired t-test; see Table 1 for detailed results; * *p* < 0.05, ** *p* < 0.01, *** *p* < 0.001) **B)** Trial-by-trial reaction time was significantly correlated to the MI for the preferred conditions across electrodes (mean Spearman’s rho = 0.051, *p* = 0.0059, two-sample t-test), but not for the non-preferred conditions (mean Spearman’s rho = 0.0031, *p* = 0.71, t-test) and the correlation was greater for the preferred than non-preferred conditions (*p* = 0.017, two-sample t-test). **C)** (left) the averaged reaction time for low MI and high MI trials in the preferred condition of the electrode (RT_low_ = 663.2 ms, RT_high_ = 681.9 ms, *p* = 0.014, permutation test); (right) the averaged reaction time for low MI and high MI trials in the non-preferred conditions of the electrode. (RT_low_ *=* 669.9 ms, RT_high_ = 669.6 ms, *p* > 0.1, permutation test) **D)** The mean correlation coefficient (Spearman’s rho) for cross-electrode correlation in the pre-stimulus MI between a pair of electrodes with the same category selectivity (left bar; mean Spearman’s rho = 0.076, *** *p* < 0.001, permutation test) versus a pair of electrodes with different category selectivity (right bar; mean Spearman’s rho = −0.0092, *p* > 0.1, permutation test).

The strength of tuning is believed to reflect the quality of the neural representation(*25*), which in turn influences the quality of perception(*26-28*). To make a connection between the aspect of endogenous activity that modulates tuning and perception, the degree to which the algorithm adjusted the classification boundary on a trial-by-trial basis was determined (the aforementioned “modulation index”; MI). The MI was correlated to reaction times for the “preferred” condition for each electrode (face trials for electrodes recording from face selective regions, word trials for electrodes recording from word selective regions, etc.), but not the “non-preferred condition (non-face trials for electrodes recording from face selective regions, etc.; Fig 2B). Furthermore, reaction times were 18.7 ms faster on average for the bottom quarter of trials than the top quarter of trials indexed by MI for the preferred condition, but was not significantly different for the non-preferred condition (Fig. 2C). These results show that the same aspect of the endogenous activity that influences tuning in a region also correlates with the trial-by-trial response time on a perceptual task in a region-by-stimulus specific manner, which supports the second hypothesis.

While the majority of the category-selective electrodes were located in the VTC, similar effects are seen in the non-VTC recordings as well. Specifically, the mean sensitivity index *d*’ = 1.00 and 0.83 with and without conditioning on prestimulus activity respectively (*t*(15) = 3.76, *p* = 0.002); the reaction times were 55.9 ms faster on average for the bottom quarter of trials than the top quarter of trials indexed by MI for the preferred condition (did not reach *p* < 0.05, but the effect is in the same direction as in VTC).

If fluctuations of endogenous activity can influence neural coding and behavior in a stimulus-specific manner then these fluctuations should be uncorrelated across regions selective for different visual stimulus categories. In particular, endogenous fluctuations could be a reflection of changes in global cognitive state, such as arousal, or general task effects, such as changes in alertness. In these cases, the MI would correlate across the brain involved in the task, regardless of category-selectivity of a particular region. However, cross-electrode correlation in MI was weakly, though statistically significantly, correlated only between electrodes that share the same category-selectivity and not for electrodes of different category-selectivity (Fig. 2D). Significantly larger correlation was seen between electrodes of the same category-selectivity than electrodes of different category-selectivity (Fig. 2D). As a result, the endogenous modulation is partially a reflection of correlated fluctuations within category-specific networks (although the effect size is weak), but it does not seem to be a reflection of non-specific processes, such as arousal or alertness, because correlations are not seen across all category selective electrodes, which supports the third hypothesis.

The results above support our three major hypotheses and shows that pre-stimulus activity can influence neural tuning and behavior in a stimulus-specific manner. A number of questions regarding the nature of the endogenous activity that influences tuning and perception remain. To evaluate the contribution of different aspects of the endogenous features, the same model was applied using different subsets of the pre-stimulus features. This analysis showed that the pre-stimulus stFP, which is dominated by the low frequency component, the pre-stimulus stBHA, which reflects the power of high frequency broadband activity, and the pre-stimulus oscillatory phase all contributed to the modulation of category tuning (Fig. 3A). The trials were then ranked by their MI and the mean and standard deviation in their pre-stimulus stFP and stBHA were compared. The bottom quarter of trials had significant lower mean and variance for both stBHA and absolute stFP during the pre-stimulus period, compared to the top quarter of trials (Fig. 3B). Given that lower MI trials corresponde to shorter RTs, the decreased pre-stimulus mean and variance may be an indication of lower endogenous noise (*29*) or fluctuations of stimulus-specific attention (*30*), which leads to shorter RTs. A further analysis into the distribution of non-zero weights in the sparse GLM suggests that the alpha/beta phases, peaked at 15 Hz, showed a consistent pattern of modulation on post-stimulus category tuning (Fig. 3C), suggesting a role for the phase of endogenous oscillations in this frequency range when visual stimuli are presented. Recent studies have shown that deployment of endogenous attention reflects neural coherence in a similar frequency range as the one seen in the current study(*31*), suggesting the prestimulus facilitation seen here may reflect an active neural process.

**Fig. 3.**
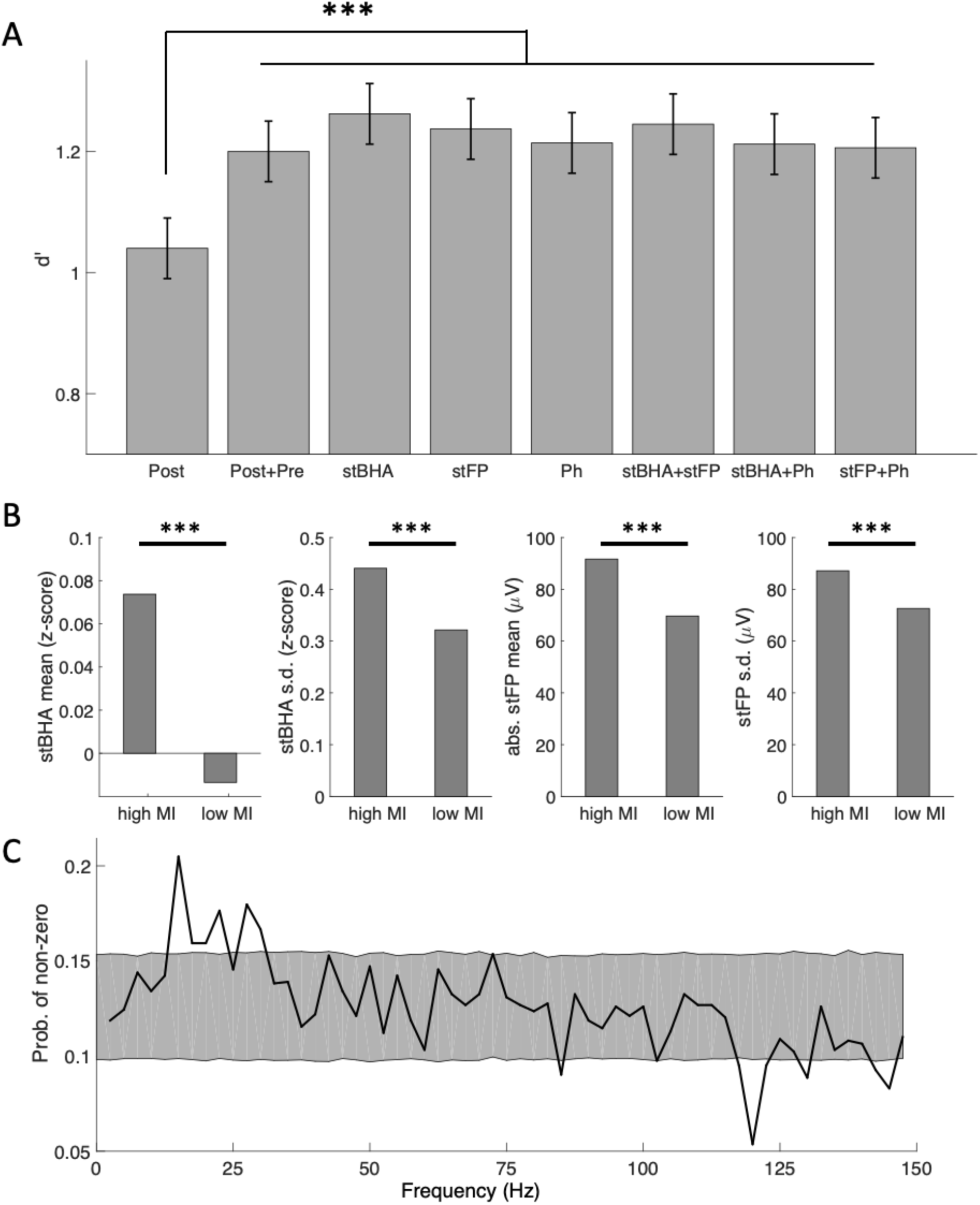
Different aspects of pre-stimulus features contributed to the modulation model. **A)** from left to right, the averaged classification *d’* across all electrodes for: (**Post**) post-stimulus features only, (**Post+Pre**) including all pre-stimulus features, (**stBHA**) including pre-stimulus stBHA features only, (**stFP**) including pre-stimulus stFP features only, (**Ph**) including pre-stimulus phase features only, (**stBHA+stFP**) including pre-stimulus stBHA and stFP features, (**stBHA+Ph**) including pre-stimulus stBHA and phase features, (**stFP+Ph**) including pre-stimulus stFP and phase features (*** *p* < 0.001, paired t-test); **B)** from left to right, 1) the averaged z-scored stBHA power, 2) the standard deviation of z-scored stBHA power, 3) the averaged absolute value of stFP, 4) the standard deviation of stFP, within [−500ms,-100ms] pre-stimulus time window for low MI and high MI trials in each electrode (*** *p* < 0.001, permutation test); **C)** the averaged empirical probability of having non-zero weights in the sparse GLM model for different pre-stimulus phase features of different frequency (shaded area: bootstrapped 95% confidence interval of being selected in the sparse GLM with random feature selection that has the same L0-norm as the current solutions);

Previous studies have shown that infra-slow fluctuations of activity, seen in “resting-state” studies, are associated with fluctuations of behavior and perception(*32-34*). If the aspect of the endogenous activity that modulates tuning and behavior seen in the present study were a reflection of these intra-slow fluctuations, there would be significant auto-correlation within each channel between consecutive trials for the MI. The auto-correlation of MI across consecutive trials for each electrode was computed, and 40 out of the 246 electrodes (∼15%) showed significant auto-correlation across trials at *p* < 0.05 uncorrected level (Fig. 4). While this is more than would be expected by chance, it is a relatively small subset of the electrodes, suggesting that there is a mix of infra-slow and transient effects in the endogenous activity, with transient effects being the dominant proportion.

**Fig. 4.**
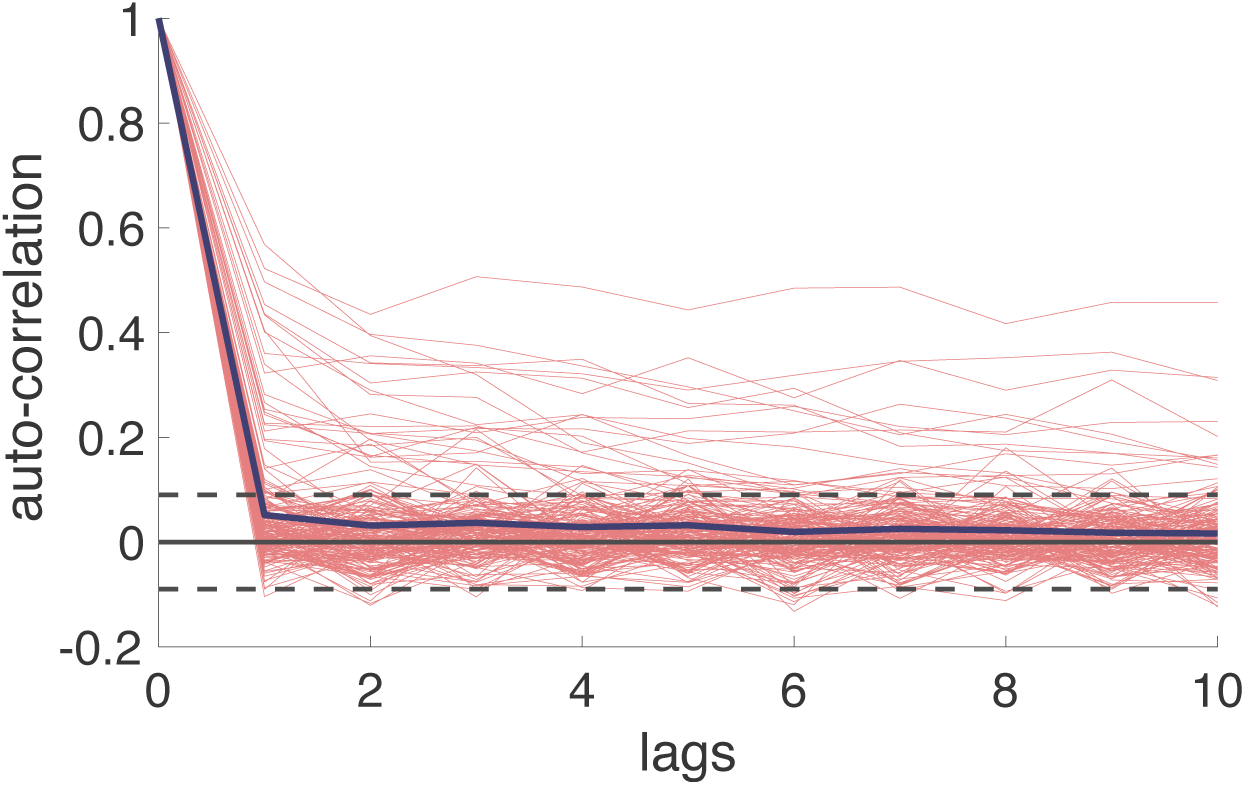
The pre-stimulus modulation effect is mostly transient. The temporal auto-correlation for MI across consecutive trials in each category-selective channel. The blue solid line indicates the average auto-correlation across all electrodes. 40/246 electrodes showed significant auto-correlation (*p* < 0.05, uncorrected). The dashed lines correspond to *p* = 0.05 threshold, uncorrected.

Taken together these results suggest a model for how endogenous states can influence neural activity to modulate the perception of specific visual stimuli. If the stimulus is presented when endogenous activity in regions selective to that type of stimulus is relatively low, as indicated by lower pre-stimulus mean and variance, and when the phases of endogenous oscillations in the alpha/beta frequency range are optimal, then neural tuning will be stronger and behavior will be facilitated. It has been suggested that endogenous fluctuations may reflect a priming-like pre-activation of a predicted stimulus(*35*), for example a prior in the Bayesian sense(*36*). However, pre-activation would likely correspond to a higher pre-stimulus response in regions that process a particular stimulus type, not lower as was seen here. Prior studies in early visual cortex in monkeys also showed that lower pre-stimulus activity is associated with improved tuning and behavior(*16, 17, 37*). The results of the present study show that these faciliatory effects can be differentially focused in circuits associated with processing specific stimulus types in higher-level visual regions and in regions outside of visual cortex in humans. Lower pre-stimulus mean and variance may reflect an optimization of the dynamic range or gain(*37*) potentially through normalization(*38*) in neural circuits responsible for processing particular stimulus types to enhance information pick-up for those stimuli(*39*). While reduced pre-stimulus activity and variance is not consistent with a priming-like prior, the results here do provide a potential foundation for endogenous activity to reflect predictive processing(*18*), though through a non-priming mechanism, such as circuit-specific optimization of processing.

Given the random stimulus presentation in the present study, facilitating one stimulus over another on a trial-by-trial basis does not provide a behavioral advantage. Therefore, it is unclear if the endogenous activity seen here reflects stochastic dynamics in brain circuits, such a fluctuations of neurotransmitter levels(*40*), or strategic processes, such as fluctuations in stimulus-specific attention or preference(*41*), that may reflect pattern detection and strategies primates adopt even when stimuli are presented randomly(*42*). In natural contexts, free viewing, and other contexts where facilitating the process of particular stimuli may be advantageous, the stimulus-specificity of endogenous optimization may reflect a prediction of the next stimulus viewed based on internal models of the environment(*35*). Active sensing in natural settings may organize the processes that underlie this optimization(*43*) and/or these active processes may synchronize to fluctuations in endogenous activity so that deployment of overt and covert attention occurs at temporally optimal times for information gathering(*44*).

Taken together, our results provide empirical support for a mechanism in which the present neural state influences the perception of sensory input in a stimulus-specific manner by modulating the tuning properties of neural circuits selective for those stimuli.

## Acknowledgments

We would like to thank the patients and staff in the epilepsy monitoring unit at the University of Pittsburgh Medical Center for their participation in this research study. We would also like to thank Shawn Walls, and Ashley C. Whiteman for their help with data collection. We thank Charles Schroeder and Matthew Smith for critically reading the manuscript and for helpful suggestions.

## Author contributions

Conceptualization, Y.L. and A.S.G.; Methodology, Y.L. and A.S.G.; Investigation, Y.L., M.J.W., R.M.R., and A.S.G..; Formal Analysis, Y.L., M.G., and A.S.G.; Writing – Original Draft, Y.L. and A.S.G.; Writing – Review & Editing, Y.L., M.G., R.M.R., and A.S.G.; Resources, R.M.R. and A.S.G.; Funding Acquisition, A.S.G.;

## Data and materials availability

Data and analysis code will be made available upon request.

## List of Supplementary Materials

Materials and Methods

Table S1 – S2

References (45 – 49)

## Supplementary Materials

### Materials and Methods

#### Subjects

The experimental protocols were approved by the Institutional Review Board of the University of Pittsburgh. Written informed consent was obtained from all participants. 30 human subjects (11 male, 19 female) underwent surgical placement of subdural electrocorticographic electrodes or stereotactic electroencephalography (together electrocorticography and stereotactic electroencephalography are referred to here as iEEG) as standard of care for seizure onset zone localization. The ages of the subjects ranged from 19 to 64 years old (mean = 38.2, SD = 11.9). None of the participants showed evidence of epileptic activity on the electrodes used in this study nor any ictal events during experimental sessions.

#### Stimuli

In each session, 180 images of faces (50% male), bodies (50% male), words, hammers, houses, and phase scrambled faces were used as visual stimuli. Each of the six categories contained 30 images, and each image was presented twice. At random, 1/3 of the time an image would be repeated, which yielded 480 independent trials in each session.

#### Paradigms

In the experiment, each image was presented for 900 ms with 900 ms inter-trial interval during which a fixation cross was presented at the center of the screen (∼ 10° x00D7; 10°of visual angle). At random, 1/3 of the time an image would be repeated, which yielded 480 independent trials in each session. Participants were instructed to press a button on a button box when an image was repeated (1-back), and their reaction time between stimulus onset and button press was recorded. Paradigms were programmed in MATLAB™ using Psychtoolbox and custom written code. All stimuli were presented on an LCD computer screen placed approximately 150 cm from participants’ heads.

#### Data preprocessing

The electrophysiological activity was recorded using iEEG electrodes at 1000 Hz. Common reference and ground electrodes were placed subdurally at a location distant from any recording electrodes, with contacts oriented toward the dura. The 60 Hz line noise was removed using a forth order Butterworth filter with 55-65 Hz stop-band. Single-trial field potential (stFP) signal was extracted by band-passing filtering the raw data between 0.2-115 Hz using a fourth order Butterworth filter to remove slow and linear drift, and high frequency noise. The stFP signal was sampled at 1000 Hz.

The single trial broadband high-frequency (stBHA) activity was defined as the mean z-scored PSD across 40-100 Hz on each trial. Specifically, power spectrum density (PSD) at 2 – 100 Hz with bin size of 2 Hz and time-step size of 10 ms was estimated for each trial using multi-taper power spectrum analysis with Hann tapers, using FieldTrip toolbox (*45*). We define the neural activity within the [−500, −100] ms interval relative to the stimulus onset as the pre-stimulus activity, and the neural activity within the [100, 500] ms interval relative to the stimulus onset as the post-stimulus activity. For each channel, the PSD at each frequency was z-scored with respect to the mean and variance of the baseline activity to correct for the power scaling over frequency at each channel. The stBHA was sampled at 100 Hz.

The pre-stimulus phase information was also extracted from each trial. Specifically, discrete time Fourier transform was applied to the raw signal in the [−500, −100] ms time interval, which had a total length of 400 points sampled at 1000 Hz. As a result, the phase information between 0-1000 Hz was extracted with a step-size of 2.5 Hz. The phases from 0 to 150 Hz were used as the pre-stimulus phase features yielding 60 phase features.

To reduce potential artifacts in the data, raw data were inspected for ictal events, and none were found during experimental recordings. Trials with maximum amplitude 5 standard deviations above the mean across all the trials were eliminated. In addition, trials with a change of more than 25 µV between consecutive sampling points were eliminated. These criteria resulted in the elimination of less than 1% of trials.

#### Electrode localization

Coregistration of grid electrodes and electrode strips was adapted from the method of Hermes, Miller, Noordmans, Vansteensel and Ramsey (*46*). Electrode contacts were segmented from high resolution post-operative CT scans of patients coregistered with anatomical MRI scans before neurosurgery and electrode implantation. The Hermes method accounts for shifts in electrode location due to the deformation of the cortex by utilizing reconstructions of the cortical surface with FreeSurfer™ software and co-registering these reconstructions with a high-resolution post-operative CT scan. SEEG electrodes were localized with Brainstorm software (*47*) using post-operative MRI co-registered with pre-operative MRI images.

#### Electrode selection

Category-selective electrodes were selected based on a 6-way classifier. Specifically, we trained a multinomial logistic regression model to classify the post-stimulus neural activity with respect to the 6 different categories from each other. The sensitivity index (*d’*) for each category was then computed as *d’* = Z(true positive rate) – Z(false positive rate), where Z(x) is the inverse function of the cumulative density function of standard normal distribution. An electrode was selected as category-selective if the maximum *d’* across all categories is greater than 0.5 (*p* < 0.01, permutation test). The selected electrode was then assigned to the category with maximum *d’*.

#### Two-stage generalized linear model (GLM)

We designed a two-stage regularized GLM (logistic regression) model to evaluate the pre-stimulus modulation on category representation.

In the first stage, logistic regression was directly applied to the post-stimulus activity to extract the critical discriminant dimensions for category classification. In other words, we solved for

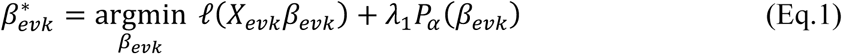

where *ℓ(x)* = −*y*^T^*x* + **1**^T^log(1 + exp(*x*)) is the cross-entropy loss for logistic regression, and 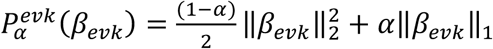 is the standard elastic-net penalty term to account for the high-dimensional settings (*48*). This results in a trial-by-trial neural metric, *X*_*evk*_ *β*_*evk*_ which corresponds to the signed distance to the classification boundary and quantifies the post-stimulus category selectivity.

In the second stage, we fixed the optimal dimension 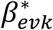 and optimized the model to modulate classification boundary along the critical discriminating directions found in the first stage, based on the pre-stimulus activity. Specifically, we solved

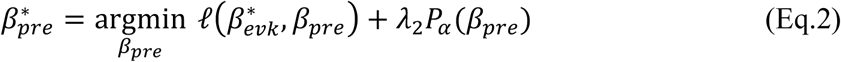

Where 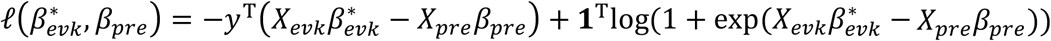, and 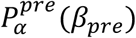 is a similar elastic-net penalty but with group structure to account for the phase features (see below for a detailed description of the penalty structure). This stage provides a neural metric *X*_*pre*_*β*_*pre*_ in pre-stimulus activity that quantifies the amount of influence from pre-stimulus activity on the post-stimulus category selectivity on a trial-by-trial basis. We defined *MI* = *X*_*pre*_*β*_*pre*_ as the pre-stimulus modulation index (MI).

We considered the neural activity within the [−500, −100] ms pre-stimulus time interval as proxy for the endogenous activity, noted as *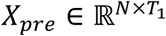*, where *N* is the number of trials and *T*_*1*_ is the number of features in the pre-stimulus time window; and we used neural activity from the [100, 500] ms time interval relative to stimulus onset as the post-stimulus evoked activity that encodes category information, noted as *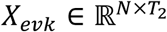*, where *T*_*2*_ is the number of features in the post-stimulus time window. The regularization parameters *λ*_1_ and *λ*_2_ were selected using cross-validation based on minimizing the deviance.

#### The (group) elastic-net penalty

For the post-stimulus part, we only considered the stFP and stBHA features, noted as *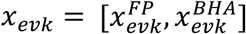*, with the corresponding weights 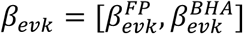, and we applied regularization term *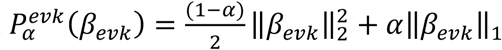*. For the pre-stimulus part, we used stFP, stBHA and phase features, noted as *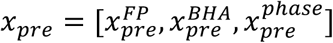*, and the corresponding weights 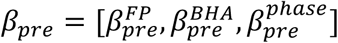. Assume that we have phase [*θ*_*1*_, …, *θ*_*k*_], where *θ* ∈ (−2*π*, 2*π*], corresponding to frequencies of interest [*f*_8_, …, *f*_K_]. To transfer the circular phase value onto the real axis in order to facilitate the *ℓ*_1_-norm penalty, we consider feature vector *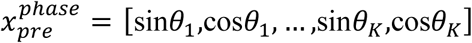*,where sin*θ*, cos*θ* ∈ [−1,1], and group lasso penalty term 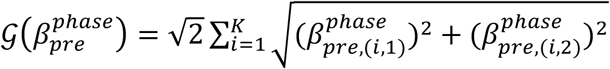, where 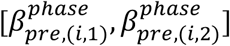 are the pair of weights corresponding to phase feature pair [sin*θ*_*i*_,cos*θ*_*i*_]. This group structure would ensure that the penalty is invariant to the overall direction of the phase, which a typical *ℓ*_1_-norm penalty would not do. As a result, the group elastic-net penalty for the pre-stimulus weights can be written as 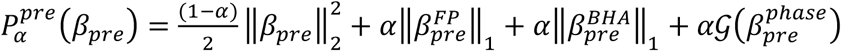.

#### Cross-electrode correlation in pre-stimulus MI

To evaluate the spatial properties of the pre-stimulus modulation effect, we computed the correlation of the single trial pre-stimulus MI between category-selective electrodes in each subject. For the *i*-th category-selective electrode, we got *MI*_*i*_ = *X*_*pre,i*_*β*_*pre,i*_ from the GLM. The cross-electrode correlation between two category-selective electrodes *i* and *j* was estimated by computing the correlation coefficient between *MI*_*i*_ and *MI*_*j*_ across all trials. To avoid confounding effect from local spatial correlation between two nearby electrodes, we only considered a pair of electrodes that were > 2cm apart from each other. For each subject, the mean cross-electrode correlation was estimated by averaging the pairwise correlation coefficients across all such pairs of category-selective electrodes.

Permutation testing was used to test for significance of the cross-electrode MI correlations (Fig. 2D). Specifically, for each permutation, we randomly shuffled the category condition of all the trials and repeat the above analysis to compute the mean cross-electrode correlation coefficients for electrodes with the same/different category selectivity. This process was repeated for 1000 times to get the histogram of the null distribution of the averaged correlation coefficient.

#### Autocorrelation in pre-stimulus MI

To evaluate the temporal properties of the pre-stimulus modulation effect, we computed the autocorrelation of the single trial pre-stimulus MI between consecutive trials with lags ranging from 1 to 20 in each category-selective electrodes. Specifically, for any given electrode, the autocorrelation with lag *k* is 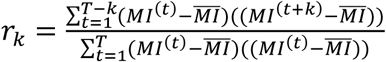. To evaluate the temporal property, we tested for the significance of the first-order autocorrelation, since it is essential for any temporal dependencies caused by slow-fluctuation in the signal. Specifically, the upper bound of the 95% confidence interval was approximately estimated as 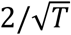 where *T* is the total number of trials.

#### Permutation test for differences based on high vs low pre-stimulus MI

Permutation test was used to test for significance of the differences in RT, pre-stimulus stFP, and pre-stimulus stHBA based on pre-stimulus MI in this study (Figure 2B, 2C, Figure 3B). In order to construct a surrogated distribution of the pre-stimulus MI, in the *i*-th permutation we generated random projection weight vector 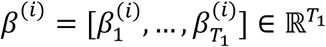, such that ‖ *β*^*i*^ ‖_0_ = ‖ *β*_*pre*_ ‖_0_. Specifically, let *n* = ‖ *β*_*pre*_ ‖_0_, we randomly drew {*p*_1_, …, *p*_*n*_} ⊂ {*p*_1_, …, *p*_1_}, and then 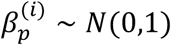 if 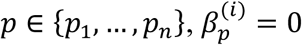 otherwise. Then we computed *MI* = *X*_*pre*_*β*^*(i)*^ and sorted the trials according to this permuted MI in order to compute the differences in RT, pre-stimulus stFP, and pre-stimulus stHBA between trials in the top quarter and trials in the bottom quarter. We repeated this process 1000 times for each electrode, and the histograms of those differences were used as the null distributions based on permuted pre-stimulus MI.

#### Classification using only pre-stimulus features

To validate that there was no discriminant information in the pre-stimulus activity, for each of the category-selective electrode, we trained a classifier using only the pre-stimulus activity. Across the 246 electrodes no significant discriminant information was presented in the pre-stimulus activity (average *d’* = 0.023; t(245) = 1.44; *p* > 0.1).

#### Concerns about category-level repetition

A possible confounding factor is the long-lasting activity from the prior trial, likely induced by the one-back task, which has been demonstrated in previous studies (*49*). This could become problematic when two consecutive trials shared the same category conditions but did not exactly repeat at the exemplar level. However, as shown in Table S1, with category-level repetitions completely removed from the trials, similar modulation effects were still found when comparing the classification accuracy with and without conditioning on the pre-stimulus activity.

#### Predicting distance to post-stimulus decision boundary

In addition to the two-stage GLM presented in the main text, a linear regression model was directly applied to evaluate the relationship between pre-stimulus activity and the absolute distance to the decision boundary in the post-stimulus discriminant model. Specifically, we solved the following linear regression problem:

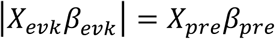

Similar to the main results presented in Figure 2 and Table 1, we found significant correlation between pre-stimulus activity and absolute distance to the decision boundary in all categories (Table S2). This suggests that the pre-stimulus activity predicts the distance to classification boundary on a trial-by-trial basis.

---

**Table S1.** The comparisons of classification results from the two-stage GLM when excluding all repeated trials with the same category as the 1-back trial.

| Category | Bodies | Faces | Words | Tools | Houses | Scrambled non-objects |
| --- | --- | --- | --- | --- | --- | --- |
| # of electrodes | 9 | 56 | 92 | 16 | 37 | 36 |
| $d'$ (evoked only) | 1.1018 | 1.5301 | 1.0847 | 0.7881 | 1.0594 | 0.8677 |
| $d'$ (evoked + endogenous) | 1.1936 | 1.6091 | 1.1904 | 0.8990 | 1.1948 | 1.0651 |
| $p$-value | 0.0908 | $1.6 \times 10^{-5}$ | $< 10^{-5}$ | $9.7 \times 10^{-4}$ | $< 10^{-5}$ | $< 10^{-5}$ |

**Table S2.** The R^2^ of the linear regression model between pre-stimulus activity and the absolute distance to the decision boundary in the post-stimulus discriminant model. (p-value estimated using the Fisher Z-transformation).

| Category | Bodies | Faces | Words | Tools | Houses | Scrambled non-objects |
| --- | --- | --- | --- | --- | --- | --- |
| # of electrodes | 9 | 56 | 92 | 16 | 37 | 36 |
| $R^2$ | 0.0717 | 0.0507 | 0.0377 | 0.0275 | 0.0361 | 0.0221 |
| $p$-value | 0.0327 | $< 10^{-5}$ | $2.78 \times 10^{-4}$ | 0.0150 | 0.0017 | 0.0678 |

